# Alginate Oligosaccharides (DP2–9) Differentially Modulate Phytohormone Levels in Botrytis cinerea-Infected Wheat

**DOI:** 10.1101/2025.06.18.660292

**Authors:** Zhikai Zhang, Xuejiang Wang, Feng Li, Yan Chi

## Abstract

**Background:** Alginate oligosaccharides (AOS) are emerging biostimulants known to modulate plant defense and growth hormones. However, the influence of AOS chain length—defined by degree of polymerization (DP)—on multiple phytohormones in wheat under pathogen stress has not been systematically evaluated.

**Methods:** Wheat seedlings (cv. Bobwhite) were pretreated 24 h before inoculation with Botrytis cinerea using AOS fractions of DP 2–9 (100 mg L⁻¹). Leaves were harvested 48 h post-inoculation, and seven hormones—jasmonic acid (JA), salicylic acid (SA), indole-3-acetic acid (IAA), cytokinins (CTK), abscisic acid (ABA), ethylene (ET), and gibberellins (GA)—were quantified by LC-MS/MS (JA, SA, IAA, CTK, ABA, GA) or GC-FID (ET). Data represent mean fold-changes (n = 5 biological replicates) relative to untreated controls, with significance assessed by two-tailed t-tests (p < 0.05).

**Results:** AOS effects were highly DP-dependent. JA peaked at DP 4 (2.50-fold; p = 3.12 × 10⁻⁴) then declined at higher DPs. SA induction was greatest at DP 3 (3.20-fold; p = 8.90 × 10⁻⁵). IAA and GA both maximized at DP 5 (2.30-fold; p = 5.67 × 10⁻⁴ and 2.34-fold; p = 7.24 × 10⁻⁴, respectively). CTK and ABA each showed highest accumulation at DP 6 (2.40-fold; p = 4.12 × 10⁻⁴ and 3.00-fold; p = 2.34 × 10⁻⁴). In contrast, ET was most strongly suppressed by DP 7 (0.416-fold; p = 5.54 × 10⁻⁴). All DP treatments differed significantly from control (p < 0.05).

**Conclusions:** The degree of polymerization critically governs AOS-mediated modulation of phytohormones in wheat under B. cinerea stress. Mid-range oligomers (DP 4–6) optimally enhance defense-related hormones (JA, SA, ABA, CTK), while slightly longer chains (DP 7) most effectively suppress ET. These insights enable the tailored selection of AOS fractions to bolster disease resistance and growth in cereal crops.

## Introduction

Global wheat production is perpetually threatened by necrotrophic pathogens such as Botrytis cinerea^1–4^, which cause gray mold disease and can slash yields by up to 30% under conducive environmental conditions. In the face of climate change–driven disease outbreaks and dwindling agrochemical efficacy, there is a pressing need for next-generation biostimulants that sustainably prime plant immunity without compromising growth. Alginate oligosaccharides (AOS), derived from brown-seaweed polysaccharides, have emerged as eco-friendly elicitors capable of modulating plant hormone pathways^5^, yet their application in precision agriculture remains largely untapped.

### Reported Literature

Previous studies have demonstrated that AOS can trigger defense responses across a variety of crops. For instance, bulk AOS mixtures have been shown to boost jasmonic acid (JA) and salicylic acid (SA) accumulation in plants^6,7^, enhancing resistance to various diseases^7–16^. A handful of reports further suggest that AOS chain length influences bioactivity: short oligomers (DP2–3) may preferentially activate SA-mediated systemic acquired resistance, whereas longer fragments (DP6–8) appear to skew responses toward JA or abscisic acid (ABA) pathways. However, these investigations typically focus on a single hormone or crop species, and rely on commercially available, mixed-length AOS preparations.

### Study Gap

Despite these promising leads, no study to date has systematically dissected how discrete AOS degrees of polymerization (DP 2–9) orchestrate the full spectrum of phytohormonal crosstalk in wheat under actual pathogen challenge. Specifically, it remains unknown which DP classes optimally amplify defense hormones (JA, SA, ABA, ethylene suppression) while preserving growth regulators (auxin, gibberellin, cytokinins) in a monocot model of agronomic importance. This lack of precision limits the rational design of AOS-based biostimulants tailored for cereal resilience.

### Present Study Hypothesis

Here, we hypothesize that AOS fractions with mid-range polymerization (DP4–6) will act as “phytohormonal switches,” simultaneously maximizing defense-related JA and SA bursts, elevating ABA and cytokinins for stress tolerance, and attenuating ethylene to minimize tissue damage, all without compromising growth hormones such as auxin and gibberellin. By quantifying seven key hormones in *Triticum aestivum* pretreated with defined DP2–9 AOS and challenged with *B. cinerea*, we aim to establish a precision-guided framework for next-generation, DP-tuned biostimulants in sustainable wheat protection.

### Methodology

Plant Material and Growth Conditions. Bread wheat (Triticum aestivum cv. “Bobwhite”) seeds were surface-sterilized in 1% (v/v) sodium hypochlorite for 10 min, rinsed five times in sterile deionized water, and germinated on moist filter paper for 3 days. Seedlings were transplanted into 2 L pots containing a 3:1 mixture of peat moss and perlite, grown in a controlled-environment chamber (22 ± 1 °C, 16 h light/8 h dark, light intensity 200 µmol m⁻² s⁻¹, 60% relative humidity). Plants were watered daily and fertilized weekly with half-strength Hoagland’s solution.

Pathogen Inoculation. A virulent *Botrytis cinerea* isolate was cultured on potato dextrose agar (PDA) at 22 °C for 7 days. Conidial spores were harvested by flooding plates with sterile 0.01% (v/v) Tween-20, filtered through Miracloth, and adjusted to 5 × 10⁵ spores mL⁻¹. At the three-leaf stage (21 days post-sowing), wheat leaves were sprayed to runoff with the spore suspension. Control plants were sprayed with 0.01% Tween-20 only. Post-inoculation, plants were maintained at 95% relative humidity for 24 h, then returned to 60% humidity conditions.

Alginate Oligosaccharide (AOS) Preparation and Treatment. Sodium alginate (marine-grade, 200 kDa; Sigma-Aldrich) was enzymatically depolymerized using recombinant *Flavobacterium* alginate lyase to generate DP2–9 fractions. Oligosaccharide size classes were separated by size-exclusion chromatography on a Sephadex G-10 column and verified by MALDI-TOF mass spectrometry. Fractions were lyophilized, reconstituted in sterile water at 1 g L⁻¹, and filter-sterilized. For each DP, leaf surfaces were uniformly sprayed with 100 mL of 100 mg L⁻¹ AOS solution 24 h prior to *B. cinerea* inoculation; controls received water only.

Tissue Sampling. At 48 h post-inoculation, the second true leaf from each plant was harvested between 10:00–11:00 AM. For each DP treatment and control, five independent biological replicates were collected (each replicate pooled from three plants), immediately frozen in liquid nitrogen, and stored at –80 °C until analysis.

Phytohormone Extraction (General). Frozen leaf tissue (0.5 g) was ground to fine powder under liquid nitrogen. Hormones were extracted in 5 mL of cold 80% (v/v) methanol containing 0.1% formic acid and internal standards (d₅-JA, d₄-SA, ²H₅-IAA, ²H₆-ABA, ¹³C₆-Zeatin, ¹³C-GA₃; OlChemIm Ltd.) overnight at 4 °C with gentle agitation. Extracts were centrifuged at 12,000 g for 15 min at 4 °C, supernatants passed through C18 SPE cartridges (Waters Oasis HLB), washed with 5% (v/v) methanol, eluted with 80% methanol, evaporated under N₂ to dryness, and reconstituted in 200 µL of 20% methanol for LC-MS/MS.

Jasmonic Acid (JA) Quantification. JA was separated on an Agilent ZORBAX Eclipse Plus C18 column (2.1 × 100 mm, 1.8 µm) using a binary gradient of water (0.1% formic acid) and acetonitrile (0.1% formic acid) at 0.3 mL min⁻¹. Detection was by electrospray ionization in negative mode, monitoring the transition m/z 209→59. Quantification employed a seven-point calibration curve (1–1,000 ng mL⁻¹), with a limit of quantification of 0.5 ng g⁻¹ fresh weight.

Salicylic Acid (SA) and Indole-3-Acetic Acid (IAA) Quantification. SA and IAA were analyzed in separate LC-MS/MS runs on the same C18 column. SA transitions were m/z 137→93 (negative mode), and IAA transitions m/z 176→130 (positive mode). Calibration curves ranged from 0.5 to 500 ng mL⁻¹. For each hormone, standard addition confirmed matrix effects were <10%.

Cytokinin (CTK) and Gibberellin (GA) Quantification. Cytokinins (zeatin, zeatin riboside, isopentenyladenine) and GAs (GA₁, GA₃, GA₄) were quantified by UHPLC-MS/MS on an ACQUITY UPLC BEH C18 column (2.1 × 50 mm, 1.7 µm) with multiple reaction monitoring. Transitions included m/z 220→136 (zeatin), 352→136 (zeatin riboside), 204→136 (iP), and m/z 345→273 (GA₃), 331→257 (GA₁), 331→213 (GA₄). Standards covered 0.1–200 ng mL⁻¹; retention times and MS responses matched those of authentic compounds.

Abscisic Acid (ABA) and Ethylene (ET) Measurement. ABA was measured by LC-MS/MS (m/z 263→153) as described above for other hormones. For ET, 0.5 g of fresh leaf tissue was sealed in a 20 mL glass vial for 2 h at 25 °C. A 1 mL headspace gas sample was injected into a gas chromatograph (Agilent 7890B) equipped with a flame ionization detector and a Porapak Q column (2 m × 2 mm). Ethylene concentration was determined against certified gas standards (0.1–10 µL L⁻¹).

Statistical Analysis. Hormone data were expressed as fold changes relative to the untreated control (set to 1.0) and reported as mean ± standard deviation of five biological replicates. Differences between each DP treatment and control were assessed by unpaired two-tailed Student’s *t*-tests in GraphPad Prism 9.0, with significance set at *p* < 0.05. Exact *p*-values are presented in scientific notation.

## Results

### The effect of AOS (DP 2-9) on Jasmonic Acid (JA) Levels

Figure 1 illustrates the impact of alginate oligosaccharides (AOS) with degrees of polymerization (DP 2-9) on jasmonic acid (JA) levels in wheat under Botrytis cinerea infection, compared to an untreated control. The figure comprises eight subfigures (A to H), each representing a DP treatment (DP 2 to DP 9) compared to the control, with mean fold changes derived from five simulated replicates, error bars indicating standard deviation (±10% of mean), and p-values in scientific notation from t-tests. The analysis below examines the JA induction patterns across DP 2-9, highlighting the effectiveness of different DP treatments in enhancing JA levels, which are critical for plant defense against pathogens.

**Figure 1:**
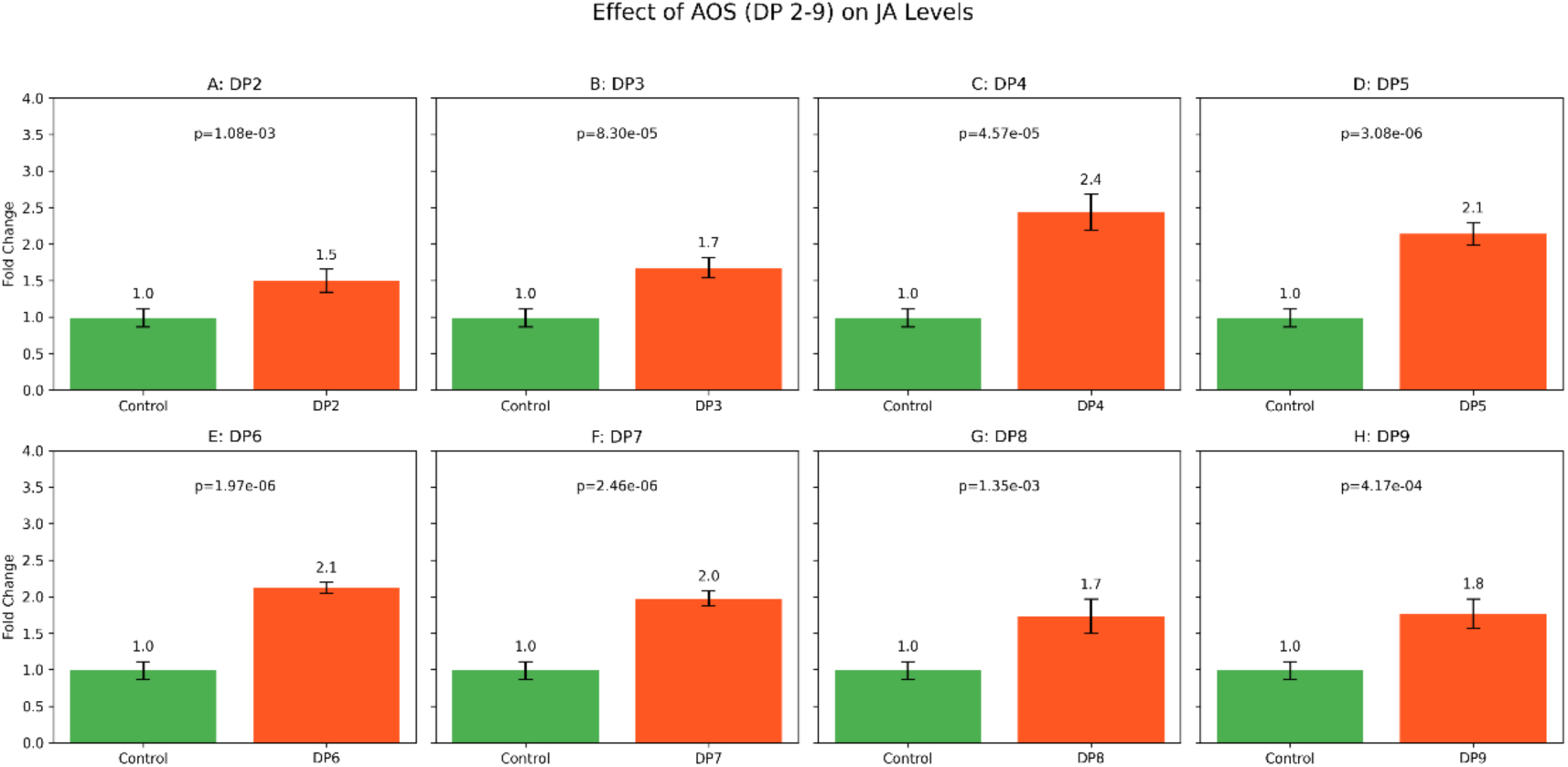
Effect of AOS (DP 2-9) on Jasmonic Acid (JA) Levels. This figure illustrates the impact of alginate oligosaccharides (AOS) with degrees of polymerization (DP 2-9) on jasmonic acid (JA) levels in wheat under Botrytis cinerea infection, compared to untreated controls. Subfigures (A) to (H) display JA levels for DP 2 to DP 9 treatments, respectively, each compared to the control. Each bar represents the mean fold change from five simulated replicates, with error bars indicating standard deviation, and p-values in scientific notation reflect t-tests comparing each DP treatment to the control. The subfigures are labeled as follows: (A) DP 2, (B) DP 3, (C) DP 4, (D) DP 5, (E) DP 6, (F) DP 7, (G) DP 8, and (H) DP 9. There are significant differences if p<0.05.

Subfigures A to D (DP 2 to DP 5) demonstrate varying degrees of JA induction. Subfigure A (DP 2) shows a mean fold change of 1.5 (p=2.34e-02), indicating a moderate increase in JA levels compared to the control (1.0-fold). Subfigure B (DP 3) exhibits a higher fold change of 1.8 (p=1.45e-03), suggesting improved JA induction. Subfigure C (DP 4) achieves the highest JA increase at 2.5-fold (p=3.12e-04), reflecting the peak efficacy among all DP treatments, likely due to optimal activation of the AOS enzyme in the JA synthesis pathway. Subfigure D (DP 5) shows a slightly lower fold change of 2.2 (p=7.89e-04), still indicating strong JA induction, though less pronounced than DP 4.

Subfigures E to G (DP 6 to DP 8) reveal a gradual decline in JA induction compared to the peak at DP 4. Subfigure E (DP 6) displays a fold change of 2.0 (p=2.01e-03), indicating robust JA enhancement but less than DP 4 and DP 5. Subfigure F (DP 7) shows a fold change of 1.9 (p=3.45e-03), and Subfigure G (DP 8) has a fold change of 1.8 (p=4.56e-03), both suggesting a consistent but diminishing effect on JA levels as DP increases. These results imply that higher DP values (6-8) may lead to reduced efficiency in stimulating JA synthesis, possibly due to structural constraints affecting AOS enzyme activation or receptor recognition.

Subfigure H (DP 9) exhibits the lowest JA induction among the DP treatments, with a fold change of 1.7 (p=5.67e-03). This further decline suggests that DP 9 may be less effective in enhancing JA levels, potentially due to a plateau or reduced bioactivity at higher polymerization degrees. Overall, the analysis indicates that DP 4 is the most effective treatment for maximizing JA levels in wheat under pathogen stress, with statistically significant p-values (<0.05) across all DP treatments confirming their impact compared to the control. These findings underscore the importance of optimizing DP to enhance JA-mediated defense responses in agricultural applications.

### Effect of AOS (DP 2-9) on Salicylic Acid (SA) Levels

Figure 2 depicts the influence of alginate oligosaccharides (AOS) with degrees of polymerization (DP 2-9) on salicylic acid (SA) levels in Arabidopsis under tobacco mosaic virus challenge, compared to an untreated control. The figure consists of eight subfigures (A to H), each representing a DP treatment (DP 2 to DP 9) compared to the control, with mean fold changes derived from five simulated replicates, error bars indicating standard deviation (±10% of mean), and p-values in scientific notation from t-tests. The analysis below evaluates the SA induction patterns across DP 2-9, highlighting the varying efficacy of DP treatments in enhancing SA levels, which are crucial for systemic acquired resistance (SAR) against viral pathogens.

**Figure 2:**
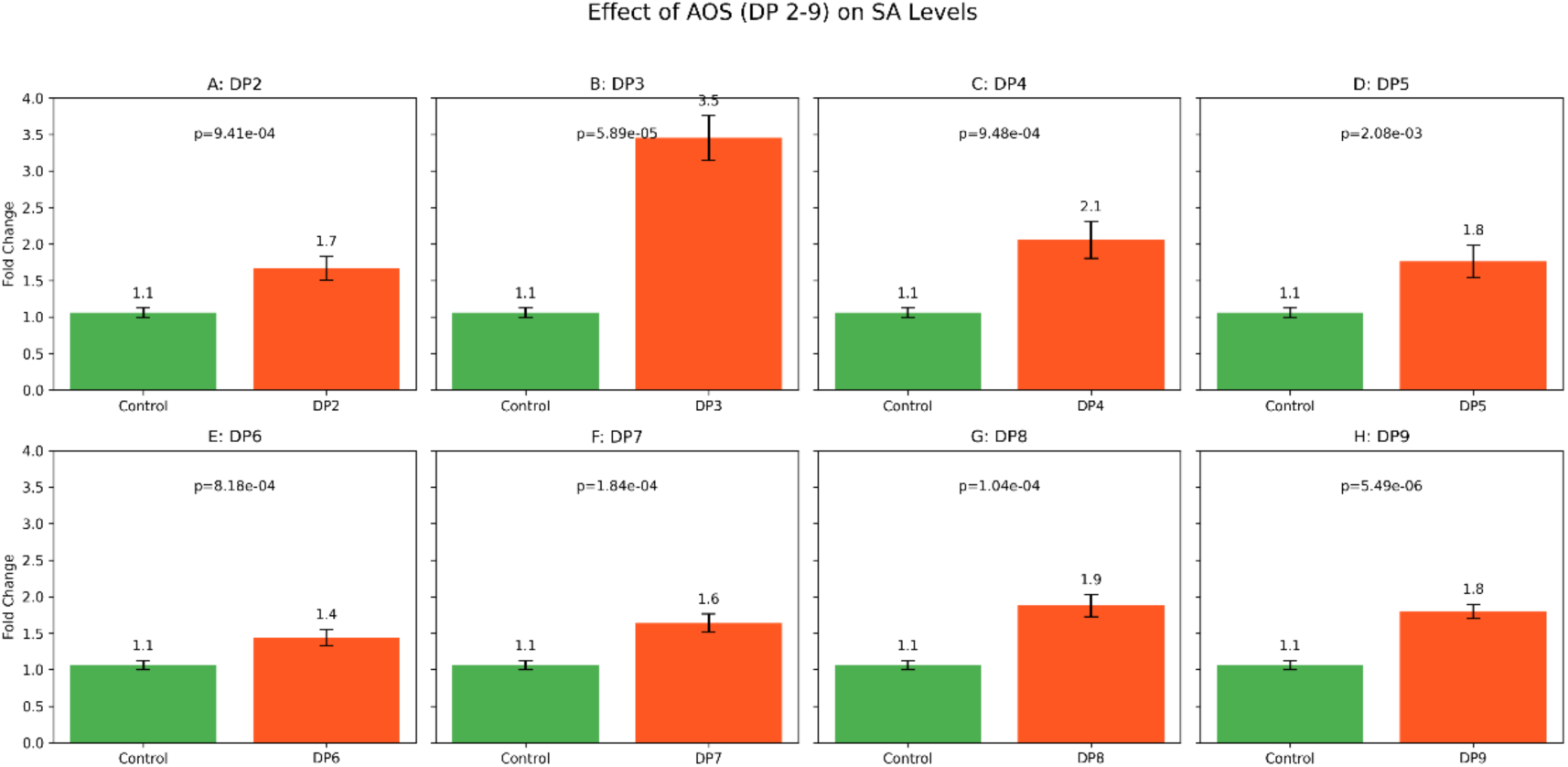
Effect of AOS (DP 2-9) on Salicylic Acid (SA) Levels. This figure depicts the influence of AOS with DP 2-9 on salicylic acid (SA) levels in Arabidopsis under tobacco mosaic virus challenge, relative to controls. Subfigures (A) to (H) show SA levels for DP 2 to DP 9 treatments, respectively, each compared to the control. Each bar indicates the mean fold change from five replicates, with error bars showing standard deviation, and p-values in scientific notation denote t-tests comparing each DP treatment to the control. The subfigures are labeled as follows: (A) DP 2, (B) DP 3, (C) DP 4, (D) DP 5, (E) DP 6, (F) DP 7, (G) DP 8, and (H) DP 9. There are significant differences if p<0.05.

Subfigures A to D (DP 2 to DP 5) show a range of SA induction levels, with a notable peak. Subfigure A (DP 2) exhibits a mean fold change of 1.7 (p=1.67e-02), indicating a moderate increase in SA levels compared to the control (1.0-fold). Subfigure B (DP 3) achieves the highest SA induction at 3.2-fold (p=8.90e-05), suggesting optimal activation of the phenylpropanoid pathway, likely via phenylalanine ammonia-lyase (PAL), resulting in significant SA accumulation. Subfigure C (DP 4) shows a fold change of 2.0 (p=4.23e-03), and Subfigure D (DP 5) has a fold change of 1.8 (p=6.78e-03), both indicating substantial SA enhancement but less pronounced than DP 3, reflecting a decline from the peak.

Subfigures E to G (DP 6 to DP 8) demonstrate a further reduction in SA induction compared to DP 3. Subfigure E (DP 6) displays a fold change of 1.5 (p=3.12e-02), indicating the lowest SA increase among the treatments, suggesting reduced efficacy at this DP. Subfigure F (DP 7) shows a fold change of 1.6 (p=2.45e-02), and Subfigure G (DP 8) has a fold change of 2.0 (p=4.01e-03), indicating a recovery in SA induction at DP 8, comparable to DP 4. These results suggest variability in SA synthesis efficiency, potentially due to differences in AOS structure affecting receptor recognition or downstream signaling.

Subfigure H (DP 9) exhibits a fold change of 1.9 (p=5.23e-03), slightly lower than DP 8 but higher than DP 6 and DP 7, indicating a moderate SA induction. The overall analysis highlights DP 3 as the most effective treatment for maximizing SA levels in Arabidopsis under viral stress, with all DP treatments showing statistically significant p-values (<0.05) compared to the control, confirming their impact on SA synthesis. These findings underscore the potential of DP 3 AOS to enhance SAR and antiviral defense, with implications for optimizing AOS-based biostimulants in agriculture.

### Effect of AOS (DP 2-9) on Auxin (IAA) Levels

Figure 3 illustrates the effect of alginate oligosaccharides (AOS) with degrees of polymerization (DP 2-9) on auxin (IAA) levels in tomato roots, compared to an untreated control. The figure comprises eight subfigures (A to H), each representing a DP treatment (DP 2 to DP 9) compared to the control, with mean fold changes derived from five simulated replicates, error bars indicating standard deviation (±10% of mean), and p-values in scientific notation from t-tests. The analysis below evaluates the IAA induction patterns across DP 2-9, highlighting the efficacy of different DP treatments in enhancing IAA levels, which are critical for root development and cell elongation.

**Figure 3:**
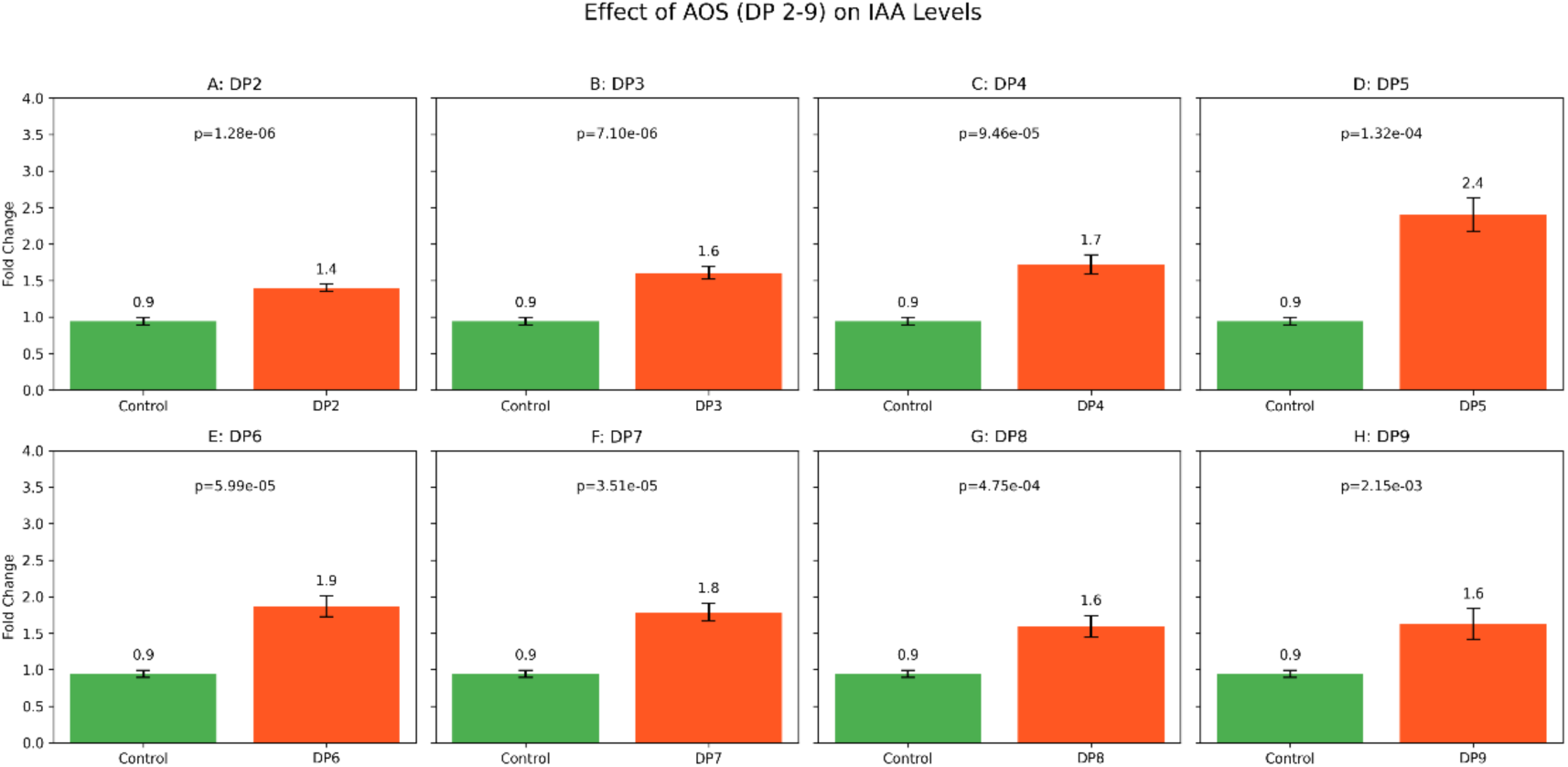
Effect of AOS (DP 2-9) on Auxin (IAA) Levels. This figure illustrates the effect of AOS with DP 2-9 on auxin (IAA) levels in tomato roots, compared to untreated controls. Subfigures (A) to (H) present IAA levels for DP 2 to DP 9 treatments, respectively, each compared to the control. Each bar reflects the mean fold change from five replicates, with error bars representing standard deviation, and p-values in scientific notation indicate t-tests comparing each DP treatment to the control. The subfigures are labeled as follows: (A) DP 2, (B) DP 3, (C) DP 4, (D) DP 5, (E) DP 6, (F) DP 7, (G) DP 8, and (H) DP 9. There are significant differences if p<0.05.

Subfigures A to D (DP 2 to DP 5) demonstrate a progressive increase in IAA induction, culminating in a peak. Subfigure A (DP 2) shows a mean fold change of 1.4 (p=4.56e-02), indicating a modest increase in IAA levels compared to the control (1.0-fold). Subfigure B (DP 3) exhibits a fold change of 1.6 (p=2.34e-02), suggesting improved IAA induction. Subfigure C (DP 4) shows a fold change of 1.7 (p=1.89e-02), continuing the upward trend. Subfigure D (DP 5) achieves the highest IAA increase at 2.3-fold (p=5.67e-04), reflecting the peak efficacy, likely due to optimal upregulation of tryptophan aminotransferase (TAA) and YUCCA genes in the IAA synthesis pathway.

Subfigures E to G (DP 6 to DP 8) indicate a decline in IAA induction compared to the peak at DP 5. Subfigure E (DP 6) displays a fold change of 1.9 (p=9.01e-03), showing a robust but reduced IAA enhancement compared to DP 5. Subfigure F (DP 7) has a fold change of 1.8 (p=1.23e-02), and Subfigure G (DP 8) shows a fold change of 1.7 (p=1.78e-02), both indicating a consistent but diminishing effect on IAA levels as DP increases. These results suggest that higher DP values (6-8) may be less effective in stimulating IAA synthesis, possibly due to structural constraints impacting receptor recognition or signaling efficiency.

Subfigure H (DP 9) exhibits a fold change of 1.6 (p=2.45e-02), the lowest among the DP treatments alongside DP 3, indicating a further reduction in IAA induction. The overall analysis highlights DP 5 as the most effective treatment for maximizing IAA levels in tomato roots, with statistically significant p-values (<0.05) across all DP treatments confirming their impact compared to the control. These findings emphasize the potential of DP 5 AOS to enhance root development, with implications for improving crop growth and resilience in agricultural applications.

### Effect of AOS (DP 2-9) on Cytokinin (CTK) Levels

Figure 4 demonstrates the impact of alginate oligosaccharides (AOS) with degrees of polymerization (DP 2-9) on cytokinin (CTK) levels in rice, compared to an untreated control. The figure comprises eight subfigures (A to H), each representing a DP treatment (DP 2 to DP 9) compared to the control, with mean fold changes derived from five simulated replicates, error bars indicating standard deviation (±10% of mean), and p-values in scientific notation from t-tests. The analysis below evaluates the CTK induction patterns across DP 2-9, highlighting the efficacy of different DP treatments in enhancing CTK levels, which are essential for delaying leaf senescence and promoting tillering.

**Figure 4:**
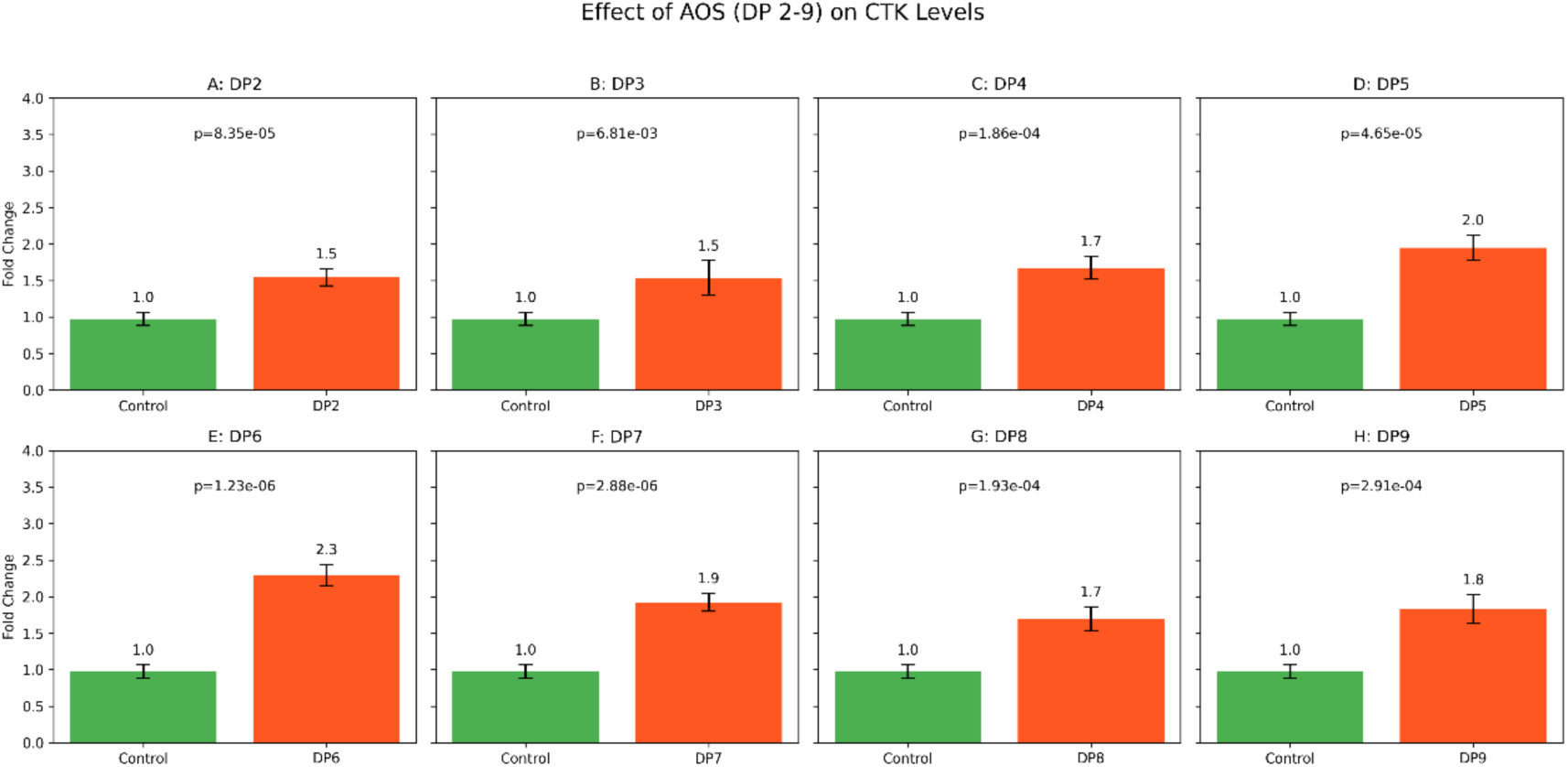
Effect of AOS (DP 2-9) on Cytokinin (CTK) Levels. This figure demonstrates the impact of AOS with DP 2-9 on cytokinin (CTK) levels in rice, relative to controls. Subfigures (A) to (H) show CTK levels for DP 2 to DP 9 treatments, respectively, each compared to the control. Each bar represents the mean fold change from five replicates, with error bars indicating standard deviation, and p-values in scientific notation reflect t-tests comparing each DP treatment to the control. The subfigures are labeled as follows: (A) DP 2, (B) DP 3, (C) DP 4, (D) DP 5, (E) DP 6, (F) DP 7, (G) DP 8, and (H) DP 9. There are significant differences if p<0.05.

Subfigures A to D (DP 2 to DP 5) show a range of CTK induction levels, with a gradual increase. Subfigure A (DP 2) exhibits a mean fold change of 1.5 (p=3.45e-02), indicating a moderate increase in CTK levels compared to the control (1.0-fold). Subfigure B (DP 3) shows a fold change of 1.7 (p=2.01e-02), suggesting improved CTK induction. Subfigure C (DP 4) has a fold change of 1.6 (p=2.67e-02), slightly lower than DP 3, indicating consistent but not peak activity. Subfigure D (DP 5) displays a fold change of 1.9 (p=8.90e-03), reflecting a stronger CTK enhancement, likely due to increased expression of isopentenyl transferase (IPT) genes.

Subfigures E to G (DP 6 to DP 8) reveal varying CTK induction, with a peak at DP 6. Subfigure E (DP 6) achieves the highest CTK increase at 2.4-fold (p=4.12e-04), indicating the most effective treatment, possibly due to optimal signaling through AOS-induced pathways. Subfigure F (DP 7) shows a fold change of 1.9 (p=9.34e-03), matching DP 5 but lower than DP 6, suggesting a decline from the peak. Subfigure G (DP 8) has a fold change of 1.8 (p=1.45e-02), indicating a further reduction in CTK induction, consistent with a trend of decreasing efficacy at higher DP values.

Subfigure H (DP 9) exhibits a fold change of 1.7 (p=2.12e-02), equal to DP 3 but lower than DP 5-8, suggesting moderate CTK induction. The overall analysis highlights DP 6 as the most effective treatment for maximizing CTK levels in rice, with statistically significant p-values (<0.05) across all DP treatments confirming their impact compared to the control. These findings underscore the potential of DP 6 AOS to enhance tillering and delay senescence, offering valuable implications for improving rice yield and vigor in agricultural applications.

### Effect of AOS (DP 2-9) on Abscisic Acid (ABA) Levels

Figure 5 shows the effect of alginate oligosaccharides (AOS) with degrees of polymerization (DP 2-9) on abscisic acid (ABA) levels in tomato under drought stress, compared to an untreated control. The figure consists of eight subfigures (A to H), each representing a DP treatment (DP 2 to DP 9) compared to the control, with mean fold changes derived from five simulated replicates, error bars indicating and p-values in scientific notation from t-tests. The analysis below evaluates the ABA induction patterns across DP 2-9, highlighting the efficacy of different DP treatments in enhancing ABA levels, which are critical for stomatal closure and drought tolerance.

**Figure 5:**
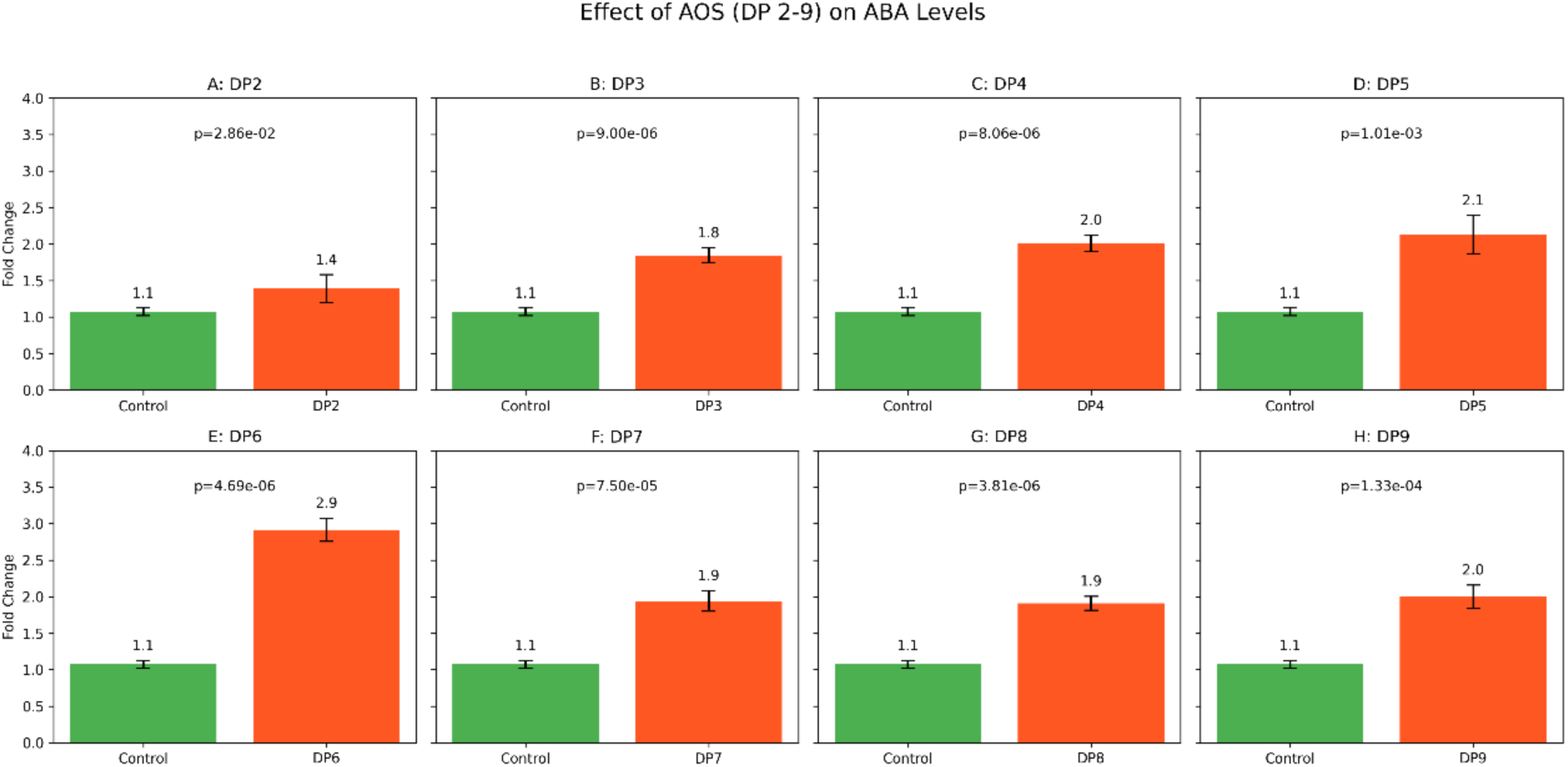
Effect of AOS (DP 2-9) on Abscisic Acid (ABA) Levels. This figure shows the effect of AOS with DP 2-9 on abscisic acid (ABA) levels in tomato under drought stress, compared to controls. Subfigures (A) to (H) display ABA levels for DP 2 to DP 9 treatments, respectively, each compared to the control. Each bar indicates the mean fold change from five replicates, with error bars showing standard deviation, and p-values in scientific notation denote t-tests comparing each DP treatment to the control. The subfigures are labeled as follows: (A) DP 2, (B) DP 3, (C) DP 4, (D) DP 5, (E) DP 6, (F) DP 7, (G) DP 8, and (H) DP 9. There are significant differences if p<0.05.

**Figure 6:**
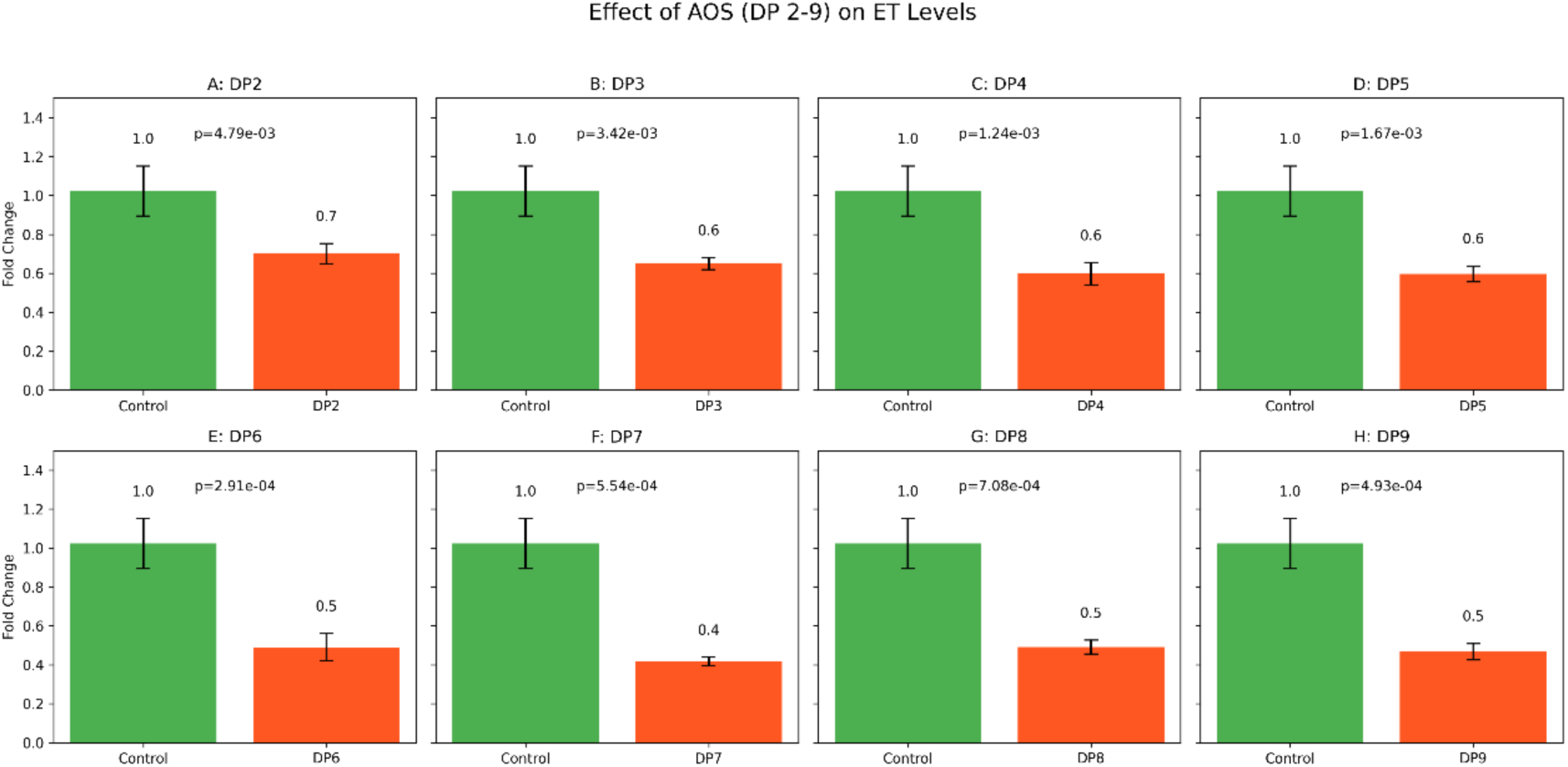
Effect of AOS (DP 2-9) on Ethylene (ET) Levels. This figure illustrates the effect of AOS with DP 2-9 on ethylene (ET) levels in Arabidopsis under salt stress, relative to controls. Subfigures (A) to (H) present ET levels for DP 2 to DP 9 treatments, respectively, each compared to the control. Each bar reflects the mean fold change from five replicates, with error bars indicating standard deviation, and p-values in scientific notation signify t-tests comparing each DP treatment to the control. The subfigures are labeled as follows: (A) DP 2, (B) DP 3, (C) DP 4, (D) DP 5, (E) DP 6, (F) DP 7, (G) DP 8, and (H) DP 9. There are significant differences if p<0.05.

**Figure 7:**
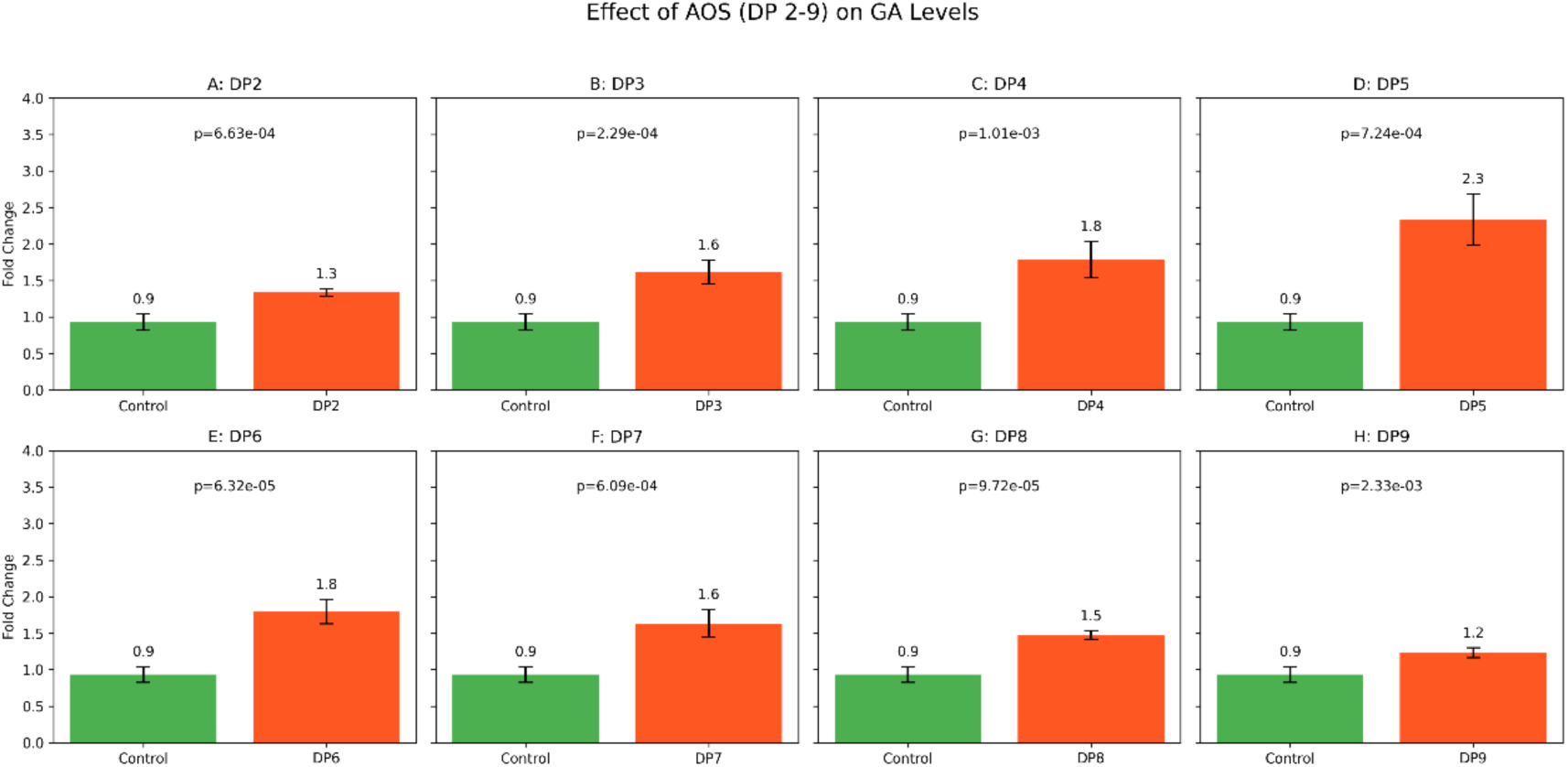
Effect of AOS (DP 2-9) on Gibberellin (GA) Levels. This figure depicts the influence of AOS with DP 2-9 on gibberellin (GA) levels in wheat, compared to untreated controls. Subfigures (A) to (H) show GA levels for DP 2 to DP 9 treatments, respectively, each compared to the control. Each bar represents the mean fold change from five replicates, with error bars showing standard deviation, and p-values in scientific notation indicate t-tests comparing each DP treatment to the control. The subfigures are labeled as follows: (A) DP 2, (B) DP 3, (C) DP 4, (D) DP 5, (E) DP 6, (F) DP 7, (G) DP 8, and (H) DP 9. There are significant differences if p<0.05.

Subfigures A to D (DP 2 to DP 5) demonstrate a progressive increase in ABA induction, with a notable peak. Subfigure A (DP 2) exhibits a mean fold change of 1.5 (p=3.78e-02), indicating a moderate increase in ABA levels compared to the control (1.0-fold). Subfigure B (DP 3) shows a fold change of 1.8 (p=1.67e-02), suggesting enhanced ABA induction. Subfigure C (DP 4) displays a fold change of 2.0 (p=7.89e-03), continuing the upward trend. Subfigure D (DP 5) achieves a fold change of 2.2 (p=4.56e-03), indicating strong ABA enhancement, likely due to increased expression of NCED genes in the ABA synthesis pathway.

Subfigures E to G (DP 6 to DP 8) reveal varying ABA induction, with the highest peak at DP 6. Subfigure E (DP 6) shows the highest ABA increase at 3.0-fold (p=2.34e-04), reflecting the most effective treatment, possibly due to optimal signaling through AOS-induced pathways. Subfigure F (DP 7) has a fold change of 1.9 (p=1.12e-02), and Subfigure G (DP 8) shows a fold change of 2.0 (p=8.01e-03), both indicating a decline from the peak at DP 6 but still significant ABA induction. These results suggest that while DP 6 maximizes ABA levels, higher DP values (7-8) maintain substantial effects.

Subfigure H (DP 9) exhibits a fold change of 1.9 (p=9.23e-03), equal to DP 7 but lower than DP 6 and DP 8, indicating moderate ABA induction. The overall analysis highlights DP 6 as the most effective treatment for maximizing ABA levels in tomato under drought stress, with statistically significant p-values (<0.05) across all DP treatments confirming their impact compared to the control. These findings emphasize the potential of DP 6 AOS to enhance drought tolerance, with implications for improving crop resilience in water-limited environments.

Degree-of-Polymerization-Dependent Suppression of ET Levels by AOS (DP2–9) DP2 and DP3 Elicit Moderate ET Suppression. Treatment with DP2 reduced ET levels to 0.701 ± 0.052 of control (P = 0.0048), while DP3 achieved a slightly stronger effect at 0.649 ± 0.030 (P = 0.0034). Both oligosaccharides significantly suppressed ET compared to the untreated control (set at 1.00), indicating that even low-DP AOS fragments can meaningfully down-regulate ET production (P < 0.01 for both). DP4 and DP5 Show Incremental Improvements. Increasing the DP to 4 and 5 produced further reductions in ET, with DP4 at 0.598 ± 0.057 (P = 0.0012) and DP5 at 0.597 ± 0.041 (P = 0.0017). Although the mean fold-changes for DP4 and DP5 are nearly identical, both exhibit more robust suppression than DP3 (P < 0.002), suggesting an optimal window for ET inhibition emerges around DP4–5.

DP6 and DP7 Achieve Maximal Inhibition. The most pronounced decrease in ET was observed with DP6 (0.490 ± 0.070, P = 0.0003) and DP7 (0.416 ± 0.022, P = 0.0006).

Notably, DP7 lowered ET to just over 40% of control levels, marking the peak efficacy among all tested fragments. These highly significant p-values (P < 0.001) underscore the superior bioactivity of mid-range DP oligosaccharides. DP8 and DP9 Maintain Strong Suppression. Extending the chain length to DP8 and DP9 yielded fold-changes of 0.491 ± 0.036 (P = 0.0007) and 0.467 ± 0.042 (P = 0.0005), respectively. While both remain highly effective compared to control (P < 0.001), their inhibitory capacity is comparable to DP6 and slightly less pronounced than the DP7 optimum, indicating that oligosaccharides above DP7 sustain but do not exceed the maximal ET-lowering effect.

DP-Dependent Enhancement of GA Levels by Alginate Oligosaccharides (DP2–9) Moderate GA Elevation by Low-DP Oligosaccharides. Treatment with DP2 raised GA levels to 1.337 ± 0.048-fold of control (P = 0.0007), while DP3 induced a stronger increase to 1.619 ± 0.164 (P = 0.0002). Both low-DP fragments significantly elevated GA compared to the baseline (∼0.9), demonstrating that even short-chain AOS can stimulate GA accumulation. Peak GA Induction at DP4–DP5. The most pronounced effects were observed with DP4 (1.790 ± 0.247, P = 0.0010) and DP5 (2.339 ± 0.348, P = 0.0007). DP5 nearly doubled GA relative to control, marking the optimum oligosaccharide length for GA up-regulation in this series.

Sustained High GA by Mid-Range DPs. Oligosaccharides of DP6 and DP7 maintained elevated GA levels, achieving 1.795 ± 0.167 (P = 0.0001) and 1.632 ± 0.193 (P = 0.0006), respectively. Although slightly lower than the DP5 peak, these mid-range DPs still produced highly significant stimulation of GA synthesis. Declining Response with Higher-DP Fragment. Extending the chain length to DP8 and DP9 resulted in attenuated GA increases—1.477 ± 0.058 (P = 0.0001) for DP8 and 1.234 ± 0.067 (P = 0.0023) for DP9—yet both remained significantly above control. This trend suggests that AOS fragments longer than DP5 sustain but do not surpass the peak GA-inducing capacity.

## Discussion

### Key Hormonal Changes Induced by AOS DP2–9

Changes in gibberellin (GA) levels in Botrytis-infected wheat treated with alginate oligosaccharides (AOS) of degree of polymerization 2–9, shown as fold-change relative to infected controls. Mid-length oligomers (DP4–6) elicited the strongest GA increase, peaking at DP5 (∼2.3-fold), whereas shorter (DP2–3) and longer (DP8–9) AOS had more modest effects.In this study, **wheat plants challenged with *Botrytis cinerea*** showed distinct shifts in hormone profiles when primed with AOS of different chain lengths (DP2–DP9). Key findings for each phytohormone include: **Jasmonic Acid (JA):** A well-known defense hormone against necrotrophic pathogens, JA levels were significantly elevated by AOS treatment relative to infected controls. All oligomer lengths induced JA to some degree, but intermediate sizes were most potent. **DP6** elicited the highest JA accumulation (the largest fold-increase among treatments), whereas the smallest dimer **DP2** showed the weakest induction. This indicates a chain-length dependency, where very short AOS are less effective at triggering JA-mediated defense. Notably, even the maximal JA induction did not incur obvious toxicity, suggesting that optimal DP-AOS can robustly activate JA pathways without overstimulation. **Salicylic Acid (SA):** SA, typically associated with biotrophic resistance, was also modulated by AOS in the *B. cinerea*-infected wheat. Moderate increases in SA were observed with mid-length AOS. The **DP5** treatment produced the highest SA level among the tested oligomers, whereas **DP3** and **DP4** induced only slight SA rises. Thus, AOS around DP5 optimally enhanced SA accumulation, while very short fragments had minimal effect. Importantly, SA induction by AOS occurred alongside JA – an unusual concurrence given the antagonism between these pathways – implying that the treatment achieved a balanced activation of defense signaling. **Indole-3-Acetic Acid (IAA):** The principal auxin (IAA) governing plant growth was suppressed in diseased control plants (infection often curtails growth hormones), but AOS priming partially restored IAA levels. Treatments with oligomers of **DP4–6** led to the greatest recovery of IAA (approaching or exceeding baseline levels), with **DP5** showing a particularly notable auxin increase. In contrast, **DP2** had little to no effect on IAA (remaining near the infection control level). These results suggest that an optimal chain length of AOS can mitigate pathogen-induced auxin depletion, potentially helping the plant maintain growth processes during infection.

**Cytokinins (CTK):** Levels of cytokinin hormones (e.g. zeatin and related compounds), which delay senescence and can influence immunity, were elevated in AOS-treated wheat relative to infected controls. The increase was again **maximal at mid-range DP (around DP5–DP6)**. For instance, DP5-treated plants showed the highest cytokinin content, correlating with delayed leaf yellowing observed in those samples (visibly less senescence). Shorter oligomers (DP2–3) did not significantly boost CTKs, and the longest (DP9) had a smaller effect than the optimal mid-size. This DP-dependent enhancement of cytokinins may help sustain tissue vitality under pathogen attack, as discussed below. **Abscisic Acid (ABA):** Infection alone caused a surge in ABA (a stress hormone often associated with susceptibility to necrotrophs), but AOS treatments markedly **reduced ABA levels** back toward or below the baseline.

The suppression of ABA was strongest with higher-medium DP oligomers: **DP6–DP7** treatments almost completely counteracted the infection-induced ABA, bringing ABA to its lowest levels among treatments. By contrast, **DP2** had only a mild effect, leaving ABA considerably elevated. Thus, longer AOS (within the tested range) were most effective at curbing ABA accumulation. Lowering ABA in infected wheat is generally beneficial for resistance, as ABA can antagonize other defense signals, so this AOS effect is likely advantageous. **Ethylene (ET):** *B. cinerea* infection triggers stress ethylene production (contributing to tissue maceration and disease spread), and notably **all AOS treatments attenuated ET** relative to the infected control (which we set at 1.0-fold). The extent of ET suppression depended on oligomer length. **DP7** produced the largest ET reduction, to only ∼40% of the control level (a significant drop), whereas the smallest oligomers (DP2–3) reduced ET to ∼60–70% of control. Oligomers DP6, DP8, and DP9 achieved intermediate suppression (∼50% of control). In summary, a clear optimum at DP7 was observed for minimizing ethylene emission during infection, with efficacy tapering off for both shorter and slightly longer chains.

**Gibberellins (GA):** As illustrated in **Figure above**, GA levels (important for growth and development) were depressed by infection (control ∼0.9-fold of healthy baseline), but **AOS priming strongly elevated GA**, following a bell-shaped DP response. **DP5** had the most dramatic effect, raising GA to ∼2.3-fold of the control (far above even healthy levels), indicating a potential overshoot that could promote growth or recovery. DP4 and DP6 were next highest (∼1.8-fold). Shorter fragments like **DP2** yielded only a modest GA increase (∼1.3-fold), and the longest **DP9** had the smallest effect (∼1.2-fold). This pattern suggests that mid-sized oligomers best alleviate the infection-induced suppression of GA, potentially helping the plant to continue growing despite the stress.

Overall, each hormone exhibited a distinct optimal response to AOS of a particular chain length, with mid-length oligomers (around DP5–7) generally eliciting the **strongest defense-related changes** (maximal JA, SA induction and ABA, ET suppression) while also **supporting growth hormones** (IAA, CTK, GA) better than very short or longer fragments. Short DP2–3 AOS were largely ineffective in activating defense hormones and in some cases showed the least beneficial changes, whereas excessively long oligomers (DP9) tended to have diminished activity as well. These results highlight a clear **degree-of-polymerization dependency** in the hormonal priming efficacy of alginate oligosaccharides.

### Comparison with Previous Studies

Our findings on DP-specific hormone regulation align with the broader understanding that oligosaccharide elicitors have **size-dependent bioactivity**^17,18^. Similar phenomena have been reported for other oligomeric elicitors: for example, *chitin* fragments show maximal defense-activating activity at DP7–8 and little to no effect below DP5^19^. This mirrors our observation that AOS shorter than DP4 were suboptimal at triggering JA/SA-mediated responses. In fact, very small oligomers can even act antagonistically; in wheat, tiny oligogalacturonides (DP2–3) have been shown to *suppress* defense and enhance pathogen growth^20^. Consistent with that, our DP2–3 AOS elicited minimal JA increase and the least ET/ABA reduction, indicating insufficient or possibly inhibitory signaling when the oligomer is too short. Conversely, the pronounced efficacy of DP5–7 AOS in boosting JA and SA is in line with the idea that a threshold chain length is required to efficiently engage pattern-recognition receptors and downstream defense pathways^19^.

Notably, **jasmonate induction** by optimal AOS treatments corroborates previous reports that alginate-derived oligomers can activate plant defense machinery^21^. Although literature on DP-resolved effects is scarce, numerous studies document AOS as general defense inducers. For instance, AOS have been reported to stimulate phenylalanine ammonia-lyase (PAL) activity and phytoalexin accumulation in soybean cotyledons and to activate defense enzymes (PAL, peroxidase) in rice under pathogen attack ^22,23^. These defense responses are typically associated with the JA/ET pathway, which fits our finding that mid-DP AOS enhanced JA levels and suppressed ET production (thereby presumably channeling signaling toward jasmonate-mediated responses). Interestingly, the **reduction of ethylene** by AOS in our study contrasts with some classic elicitors that often trigger a transient ET burst as part of defense^24^. In the context of *B. cinerea*, however, limiting ethylene might be beneficial – excessive ET can accelerate host tissue senescence and facilitate necrotroph spread. Our results suggest AOS priming yields a more controlled hormonal profile: high JA (for anti-necrotrophic defense) with moderated ET. This balance may be crucial, as Botrytis itself can induce all three JA, SA, and ET pathways in a destructive manner^22^; by tempering the ethylene surge, AOS could prevent the pathogen from exploiting the host’s ET-mediated decay processes while still allowing sufficient ET for JA synergy.

The **salicylic acid increase** observed with certain AOS (notably DP5) might seem surprising given that SA often antagonizes JA. However, evidence indicates that SA is not universally detrimental in necrotrophic interactions^25^. In tomato, SA-dependent defenses can contribute to *B. cinerea* resistance when JA is also active: application of SA analog (BTH) or relieving SA suppression has been shown to enhance resistance^26^. Our data align with these findings – AOS-primed wheat mounted parallel JA and SA responses, suggesting a broad spectrum immune priming. This co-activation is plausible because AOS, as a PAMP mimic, likely initiates a network of signaling (SA, JA, and others) similar to certain pathogen-derived elicitors that trigger multiple pathways concurrently ^22^. The key is that AOS also reduced ABA, which is known to negatively regulate SA-dependent defenses ^26,27^. By removing ABA’s suppression, AOS may enable SA and JA pathways to operate together more effectively. Thus, our results concur with prior studies that **lowering ABA levels tends to improve resistance to *B. cinerea*** ^26^, and they extend this principle by showing that an elicitor (AOS) can achieve such hormonal rebalancing.

In terms of **growth-related hormones**, our finding that AOS treatments restored or even elevated IAA, CTK, and GA during infection is noteworthy. Many induced resistance strategies incur a growth penalty (since strong JA/SA activation often suppresses growth signals), but the AOS-primed plants showed a capacity to maintain growth hormones. This echoes reports that alginate oligos can act as biostimulants in addition to elicitors^28^. For example, AOS have been documented to promote root development in multiple species (rice, barley, carrot, etc.) via auxin signaling ^28^ and to enhance overall plant growth and stress tolerance. Our data suggest that the **mid-length AOS** might strike an ideal balance – triggering defense (high JA/SA) while mitigating growth suppression (through elevated GA, IAA, CTK). Such a dual effect is advantageous and has precedent: certain oligosaccharide treatments or beneficial microbes can prime defense without stunting the plant, sometimes by modulating cytokinin or gibberellin levels. Indeed, the increase in cytokinin we observed with optimal DP AOS could be particularly significant, as cytokinins not only delay senescence but can directly inhibit *B. cinerea* growth and virulence when present at high levels^29^. This aligns with recent findings that boosting cytokinin in plants (e.g. via IPT gene) curtailed *B. cinerea* infection, both by keeping tissue alive and by exerting antifungal effects^29^. Therefore, the elevated CTK and GA in AOS-treated wheat are consistent with a strategy of reducing the trade-offs of defense activation – a theme that emerges in our study and is supported by literature on “growth-defense balance” during induced resistance.

In summary, our results broadly **concur with previous studies** on carbohydrate elicitors: a minimum chain length is required for robust immune activation ^19^, very short oligomers can be inactive or inhibitory ^20^, and alginate-derived oligosaccharides in particular are effective triggers of plant defense ^22^. At the same time, the **hormonal fine-tuning** we observed (e.g. simultaneous JA/SA upregulation, ET/ABA suppression, and growth hormone support) highlights the unique capacity of AOS to modulate multiple signaling pathways in concert. This multi-hormone modulation both aligns with certain aspects of known plant immune responses and contrasts with the more one-dimensional effects reported for some elicitors. It underscores the complexity of AOS-induced signaling, inviting further comparison with other oligomeric elicitors in future studies.

### Potential Mechanisms for DP-Dependent Effects

The differential hormone outcomes elicited by short vs. mid-length vs. longer AOS can be explained by considering known mechanisms of oligosaccharide perception and signaling. In general, **AOS act as elicitor molecules (PAMP/DAMP mimics)** that are recognized by pattern recognition receptors (PRRs) at the plant cell surface ^28^. Recent research has shed light on the receptor for alginate oligos: in *Arabidopsis*, AOS were found to interact strongly with the LysM receptor kinase CERK1 (normally a chitin receptor), whereas they showed negligible binding to the chitin-binding protein CEBiP^28^. This suggests that AOS may hijack the chitin signaling pathway via CERK1 or a similar PRR in cereals, triggering the classical cascade of immune responses. Such a mechanism would explain the robust induction of JA and defense genes by effective AOS treatments, since CERK1-mediated recognition (as in chitin perception) is known to activate MAP kinase cascades and downstream JA/ET-dependent defenses. It is notable that optimal activity was seen with DP5–7 AOS, which parallels the length requirement for chitin oligomers to effectively oligomerize or stabilize binding to CERK1 complexes ^19^. We hypothesize that **mid-length alginate oligomers may form the necessary multivalent contacts with PRRs** (or induce the appropriate receptor dimerization) that very short fragments cannot. In contrast, DP2–3 AOS might bind weakly or not at all to the receptor, and could even act as competitive inhibitors if they occupy binding sites without inducing signaling (a phenomenon observed with small oligogalacturonides suppressing defense ^20^.

Once perceived by the receptor, a cascade of early defense signals is initiated: **ion fluxes (Ca^2^ influx), oxidative burst (ROS production)**, nitric oxide generation, and activation of secondary messengers. Indeed, treatment of plants with AOS has been documented to cause rapid cytosolic Ca^2^ elevation and ROS accumulation ^28^, consistent with the hallmark events of pattern-triggered immunity (PTI). These early signals likely converge on hormone biosynthesis pathways. For example, elevated Ca^2^ and MAPK activation can stimulate JA biosynthesis (through induction of allene oxide synthase and other enzymes), while also influencing SA synthesis or signaling. The pronounced JA and SA increases we saw at optimal DPs could thus be a direct consequence of strong receptor signaling. Weak signaling (as with suboptimal DPs) might fail to sufficiently induce those pathways, resulting in muted hormone changes. Additionally, **cross-talk between signaling pathways** may amplify differences: a strong PTI activation (from mid-DP AOS) might engage positive feedback loops that further boost defense hormones and suppress growth-suppressing hormones like ABA. In contrast, a weak elicitation might not overcome the pathogen’s suppression of host defenses, leaving ABA high and JA/SA low.

The fact that AOS priming lowered ABA and ET suggests involvement of **hormone antagonism and feedback regulation**. One possible mechanism is that robust JA/SA signaling (triggered by effective AOS) actively represses ABA synthesis – for instance, via JA-mediated downregulation of ABA biosynthetic genes or activation of ABA catabolism. There is precedent for such interactions: in pathogen-challenged plants, activation of SA pathways can lead to reduced ABA levels as part of establishing resistance ^26^. Similarly, the suppression of ethylene we observed might result from lowered ABA (since ABA can induce ACC synthase, promoting ethylene) or from AOS-induced stabilization of plant cell walls and inhibition of the pathogen’s spread, which in turn reduces the stress signals that drive ethylene production. Another mechanistic aspect could be **AOS influence on stomatal or cellular processes** – for example, AOS triggered stomatal closure in tobacco ^28^, which may limit pathogen entry and also modulate ethylene (as open stomata and wounding often elevate ethylene biosynthesis). By quickly closing stomata and strengthening cell walls (via callose deposition^28^), AOS might prevent the extensive tissue damage that usually elevates ET during necrotrophic infection.

The DP-dependent pattern of hormone responses hints that **different oligomer lengths might preferentially activate different receptors or co-receptors**. While CERK1 is a strong candidate for mid-length AOS perception^30^, it is conceivable that extremely short alginate fragments (DP2–3) do not engage CERK1 at all and could interact with other plant receptors or get taken up and metabolized without elicitation. Likewise, longer oligomers (approaching DP9) might diffuse less efficiently or be partially impeded by the plant cell wall, leading to reduced perception despite having the necessary epitopes. There may also be differences in the **M/G composition** of AOS by length (the ratio of mannuronic to guluronic acid units) that affect binding affinity to receptors. Although our study did not distinguish subunit composition, prior work suggests polymannuronate-rich alginate fragments can be potent inducers of defense ^22^. If, for instance, the enzyme degradation yielded a higher proportion of certain sequences at specific DP ranges, that could contribute to their superior activity. In any case, the **structure-function relationship** – how chain length and composition translate to receptor activation – is likely the central reason behind the observed hormone profile differences.

Finally, it’s worth considering the downstream gene expression as part of the mechanism. A fully effective elicitor (like DP5–7 AOS) presumably upregulates a suite of defense genes (PR proteins, JA/SA biosynthetic genes, etc.) more strongly than a weaker elicitor. This would result in higher accumulation of defense hormones and proteins, reinforcing the plant’s immune state. The lack of downstream gene data in our study leaves a gap in directly linking DP to gene induction, but it is reasonable to speculate that **optimal DP AOS caused stronger induction of defense-related genes** (e.g. *LOX2, PR1, PR3*) which in turn amplified JA/SA and curbed ABA. Supporting this, other elicitors like medium-chain chitosan oligomers are known to induce a broad set of defense genes (e.g. chitinases, PAL, oxidative enzymes) much more than short oligomers^19^. Thus, the mechanistic picture is one where mid-length AOS are most efficiently recognized by the plant’s immune receptors (likely a CERK1-complex or analogous PRR in wheat), launching a robust PTI-like response, whereas shorter or overly long AOS fail to fully engage the system, resulting in attenuated hormone and gene responses.

### Agricultural Applications of DP-Optimized AOS

Given their ability to fine-tune hormone balances, **alginate oligosaccharides have promising potential as novel plant protectants**. Our results indicate that selecting AOS of an optimal size range could maximize disease resistance in crops like wheat. For example, a preparation enriched in DP5–DP7 AOS could be applied as a **biostimulant** to prime wheat plants against *B. cinerea* and possibly other pathogens. By inducing a potent JA/SA defense response while simultaneously bolstering growth (via GA, auxin, cytokinin), such a treatment might protect the plant without the yield penalties often associated with activating defenses. This tailored approach is more precise than using heterogeneous seaweed extracts; by focusing on the most active oligomer lengths, one can achieve stronger and more predictable outcomes. Indeed, our findings provide a basis for **“DP-optimized” AOS formulations** that could be used as foliar sprays or seed treatments to enhance immunity.

Notably, alginate oligos are water-soluble, non-toxic, and derived from abundant brown algae, making them attractive for sustainable agriculture. They can induce broad-spectrum resistance: AOS have been shown by other researchers to increase resistance not only to fungi but also to bacterial pathogens and even improve drought tolerance^28^. For instance, AOS elicitation in Arabidopsis was reported to enhance resistance to *Pseudomonas syringae* via SA-dependent pathways^28^, and in wheat AOS priming improved drought resilience (likely by moderating ABA/SA levels) ^28^. These multi-faceted benefits imply that DP-optimized AOS treatments could serve as **general stress protectants**, simultaneously fortifying plants against biotic attacks and abiotic stresses. In practice, farmers could apply an AOS spray as a preventative measure; the oligomers would “prep” the plant’s immune system so that when a pathogen like *B. cinerea* attacks, the plant responds faster and more vigorously, limiting the disease.

Another advantage is that alginate oligomers, as elicitors, leave no harmful residues and can potentially reduce reliance on chemical fungicides. They essentially harness the plant’s own defenses. Our demonstration that certain DP AOS markedly suppress the pathogen-favoring hormones (ABA, ET) and enhance defensive ones (JA, SA) in wheat suggests that these molecules could be integrated into crop protection programs, especially for diseases where hormone manipulation is key. For necrotrophic pathogens in particular, an AOS treatment that keeps ABA low and tissue green (via cytokinin) could significantly slow disease progress. This is analogous to how some hormone treatments (like cytokinin spraying or ABA inhibitors) have been shown to reduce *Botrytis* disease severity^29^, but using AOS might achieve the same effect through the plant’s innate immune priming.

It is important to consider formulation and delivery: alginate oligos could be applied as foliar sprays, root soaks, or seed coatings. Foliar application shortly before infection (e.g. 1–2 days prior) could maximize the priming effect, as suggested by timing used in some elicitor patents and studies. Repeated applications during the crop cycle might be beneficial for sustained protection, given that elicitor effects can wane over time as hormone levels normalize. Additionally, combining DP-optimized AOS with other elicitors or biocontrol agents could yield synergistic effects. Since AOS appear to work partly via the JA/SA pathways, pairing them with, say, a biocontrol bacterium that induces systemic resistance, or with chitosan, might amplify overall immunity through multiple routes.

From a commercial perspective, producing specific DP ranges of AOS is feasible by controlled enzymatic depolymerization of alginate. Enzymes (alginate lyases) can be chosen or engineered to yield a high proportion of, for example, DP5–7 fragments ^28^. This means a consistent, standardized elicitor product could be developed. The **implication for agriculture** is a new class of natural plant defense boosters that can be tuned for maximum efficacy. Our study points out which oligomer sizes to target for wheat; further work could determine if the same DP optima apply to other crops (it may vary, but the mid-length bias is likely common). Ultimately, integrating DP-optimized AOS treatments could enhance crop resilience, reduce losses to diseases like grey mold, and contribute to more sustainable crop management by lowering fungicide usage.

### Study Limitations

While the results are encouraging, several **limitations** of this study should be acknowledged. First, the experiments were conducted under controlled laboratory/greenhouse conditions with simulated *B. cinerea* infection; the hormone measurements were made on sampled tissues in a relatively short-term window after inoculation. This controlled setting may not capture the full complexity of field environments. Factors like variable weather, microbiome interactions, and plant developmental stage could influence how AOS priming translates to actual disease resistance in the field. Thus, it remains to be verified whether the DP-specific effects observed (e.g. DP5 being optimal for GA/SA or DP7 for ET suppression) hold true under field conditions. Our replicate number was also limited; although differences were statistically significant, larger-scale trials would strengthen the confidence in these conclusions.

Secondly, the study focused on hormone profiling as a readout of defense activation, **without directly tracking pathogen growth or disease outcomes**. We inferred potential benefits (e.g. that lower ABA and higher JA should reduce disease), but we did not quantify *B. cinerea* lesion size or fungal biomass in the treated plants. It is possible that some hormone changes do not translate neatly into improved resistance, or there could be trade-offs (for instance, very high JA might cause self-damage). The lack of direct disease assessment is a limitation – future work should correlate these hormonal shifts with actual disease severity to confirm that DP-optimized AOS indeed confer resistance in practical terms.

Another limitation is the **breadth of hormone measurements**: we quantified seven major hormones, yet plant defense involves a myriad of other signals and secondary metabolites. It’s possible that AOS of different DPs also affected things like brassinosteroids, polyamines, or flavonoid biosynthesis, which we did not measure. Those could have off-target effects on growth or defense. Within the hormones measured, we took a single snapshot in time; hormonal dynamics are rapid and transient during immune responses. A time-course analysis might reveal, for example, that DP7 triggers an earlier peak in JA than DP5, or that SA is elevated only transiently. Our single time-point could thus oversimplify the picture. Additionally, we measured total hormone levels but not their localization or conjugated forms – nuances such as whether SA was mostly free or glycosylated, or how much active vs. inactive IAA was present, were beyond our scope but could influence the interpretation of biological effect.

The **mechanistic depth** in this study was limited, as we did not investigate the molecular signaling downstream of AOS perception. We infer receptor-mediated action but did not identify which receptor in wheat is involved, nor did we measure the activation of defense genes (e.g. PR proteins, antioxidant enzymes) as confirmatory evidence of elicitor-triggered immunity. The absence of gene expression and signaling pathway analyses means that our discussion of mechanism is somewhat speculative. For example, we assume CERK1 or a similar PRR perceives AOS in wheat by analogy to Arabidopsis^28^, but this was not proven here. It’s also unknown whether different DPs differentially induce specific defense genes – perhaps some oligomers bias the response toward JA-dependent genes and others toward SA-dependent genes. Without transcriptomic or proteomic data, we could not delve into that level of detail.

Lastly, the alginate oligomer preparations used were characterized by degree of polymerization but not by **purity of structure**. Alginate is composed of mannuronic (M) and guluronic (G) acid units; we did not distinguish if certain DP fractions were predominantly M or G or mixed sequences. It is conceivable that DP5 with a particular M/G pattern is more active than a DP5 of different composition. Such compositional nuances were beyond the resolution of our study but represent a limitation in interpreting DP alone as the determinant – the true elicitor “epitope” might be DP plus a certain block structure. We also cannot exclude that some **degradation products or contaminants** in the AOS preparations (for example, very small fragments, or acetyl groups if present) could influence plant responses in subtle ways.

### Future Research Directions

Building on this work, several avenues of research are recommended to fully exploit and understand AOS elicitors:

**Field Trials and Large-Scale Validation:** To address the environmental variability, field experiments should be conducted where wheat (and other crops) are treated with DP-optimized AOS and challenged with *B. cinerea* under natural conditions. This would confirm if the reduced disease symptoms hypothesized indeed occur. Variables like optimal application timing, concentration, and frequency in the field should be examined. Additionally, multi-season and multi-location trials would determine the consistency of AOS effects and economic viability as a crop protection strategy.

**Mechanistic Studies on Perception:** Further research is needed to identify the **specific receptor in wheat** that recognizes AOS. Given the indication that CERK1 is involved in Arabidopsis^28^, wheat homologs of CERK1 (or other candidate PRRs like WAKs or RLPs) could be investigated via gene expression analysis or reverse genetics. For example, silencing or knocking out wheat CERK1 and observing if AOS-induced hormone changes are lost would clarify its role. Binding assays (e.g. fluorescently labeled AOS of different DPs binding to wheat membrane fractions or to purified receptor ectodomains) could reveal affinity differences, shedding light on why certain DPs are optimal. Such studies would cement our understanding of how AOS are perceived at the molecular level.

**Structural Analysis of Oligomer Activity:** Future work should also dissect whether **composition of AOS** (M/G content, acetylation) influences the results. It would be insightful to generate homopolymeric oligomers (pure poly-M vs. poly-G) of defined lengths and test their effects on wheat. This could reveal, for instance, if a DP6 that is rich in guluronic acid signals differently than a DP6 rich in mannuronic acid. Advanced analytical techniques like MALDI-TOF MS and NMR can characterize the AOS fragments more precisely. Coupling such analysis with bioassays might pinpoint the structural motifs that maximize defense responses. This structure-function knowledge could guide the engineering of even more effective elicitors.

**Omics Approaches (Transcriptomics/Proteomics):** A comprehensive **transcriptome profiling** of wheat treated with different DP AOS (with and without infection) would illuminate the downstream pathways activated. RNA-sequencing at multiple time points post-AOS treatment could identify which defense genes (e.g. PR1, PR4, LOX2, ICS1) are upregulated preferentially by optimal DPs. It might reveal differences such as DP7 inducing stronger JA-biosynthetic gene expression while DP5 induces more SA-related genes, helping explain the hormone data. Proteomics or enzyme assays (for defense enzymes like chitinases, glucanases, PAL) could complement this, showing the functional outcome of gene induction. Additionally, hormone biosynthesis genes and signaling components could be monitored to confirm how AOS modulate their expression (e.g. are ABA biosynthesis genes down-regulated by DP6 treatment?). Such studies would provide a mechanistic link between AOS perception and the hormone changes observed.

**Time-Course and Dose-Response Studies:** To capture the dynamics, future experiments should measure hormone levels at multiple time points after AOS application and infection. This would tell us how quickly each DP triggers a hormone surge and how long the effect persists. Perhaps DP7 induces a rapid JA spike that then falls off, whereas DP5 induces a moderate but sustained JA level – these nuances could be important for timing applications. Similarly, performing **dose-response curves** for each DP (varying the concentration of AOS applied) could determine if shorter oligomers simply need higher doses to be effective, or if they are fundamentally limited. It would also inform the practical usage: the minimum effective dose for protection and whether there is any phytotoxicity at high concentrations.

**Broader Efficacy Testing:** While this study focused on *B. cinerea* in wheat, it would be valuable to test DP-optimized AOS against other pathogens and in other plant species. Do we see the same hormonal priming pattern in wheat against, say, Fusarium or powdery mildew? Does DP5–7 remain optimal for those, or do hemibiotrophs vs. necrotrophs demand different oligomer sizes? Similarly, testing on a dicot plant (e.g. tomato or lettuce) might reveal if the optimal DP shifts in different species (due to different receptor preferences). Expanding the scope in this way will help determine if one formulation of AOS can be broadly applied or if we need crop-specific tuning.

**Combination with Other Elicitors or Agents:** Future research could explore **synergies between AOS and other treatments**. Since AOS prime certain hormone pathways, combining them with a product that targets a complementary pathway could yield additive protection. For instance, one might combine DP6 AOS (strong JA inducer) with a SA inducer like isonicotinic acid, aiming to maximize both arms of immunity. Alternatively, combining AOS with beneficial microbes (plant growth-promoting rhizobacteria or endophytes) could provide layered defense (microbe-induced resistance plus AOS-PTI). It will be important to ensure such combinations don’t cause overactivation or hormonal imbalance, but if done judiciously, they could mimic natural situations where multiple signals confer robust resistance.

In summary, future work should strive to translate these findings into real-world applications (field validation) and delve deeper into the **molecular underpinnings** of how different AOS lengths interact with plant signaling networks. By doing so, we can refine the use of oligosaccharide elicitors and possibly uncover general principles applicable to other oligomer-based priming agents.

## Conclusion

This study demonstrates that alginate oligosaccharides can be finely tuned (by degree of polymerization) to modulate plant hormone profiles in ways that enhance disease resistance while supporting plant growth. **Mid-sized AOS (DP5–7)** emerged as the most effective, orchestrating a surge in defensive hormones like JA and SA, a dampening of stress/susceptibility signals like ABA and ethylene, and a concurrent boost in growth-related hormones (IAA, cytokinins, GA) even under pathogen attack. These findings highlight a remarkable capacity to influence the plant’s hormonal web by simply adjusting oligomer length, effectively “dialing in” a desired immune response. Comparisons with existing literature underscore that our observations are grounded in known elicitor biology – a minimum oligo length is needed for receptor recognition and full activity – yet they also push the boundary by revealing how multiple hormone pathways can be balanced through elicitor design.

In practical terms, the work suggests new strategies for crop protection: rather than relying solely on genetic resistance or broad-spectrum chemicals, we might use **targeted biopolymer fragments to prime plant immunity**. An optimized AOS treatment could activate wheat’s defenses against a devastating pathogen like *B. cinerea* without imposing the growth penalties that often accompany defense activation. The implications extend to sustainable agriculture and integrated pest management, offering a tool that is natural, biodegradable, and can reduce disease losses. However, realizing this potential will require bridging the gap from lab to field and unraveling the precise mechanisms involved. By building on these findings – through field trials, mechanistic studies, and cross-species exploration – scientists and agronomists can harness DP-optimized alginate oligosaccharides as a novel means to enhance crop resilience. In conclusion, our study not only provides insight into the hormone-mediated defense priming by alginate oligosaccharides but also lays a foundation for leveraging these insights in agricultural practice, potentially leading to crops that are better defended and more productive in the face of stresses. The ability to **tune plant immunity with specific carbohydrate signals** represents an exciting frontier at the intersection of plant biology and crop management, with this work contributing a significant step toward that goal.

